# Transcriptome analysis of three *Agave* fiber-producing cultivars suitable for biochemicals and biofuels production in semiarid regions

**DOI:** 10.1101/2020.06.03.132837

**Authors:** Fabio Trigo Raya, Marina Pupke Marone, Lucas Miguel Carvalho, Sarita Candida Rabelo, Maiki Soares de Paula, Maria Fernanda Zaneli Campanari, Luciano Freschi, Juliana Lischka Sampaio Mayer, Odilon Reny Ribeiro Ferreira Silva, Piotr Mieczkowski, Marcelo Falsarella Carazzolle, Gonçalo Amarante Guimarães Pereira

## Abstract

- Agaves, which have been grown commercially for fiber or alcoholic beverages, are emerging as a candidate crop for biochemicals and biofuels production in semiarid regions because of their high productivity in low rainfall areas, drought tolerance, and low lignin content.
- In this work, we present the transcriptomic atlas of *Agave sisalana*, *Agave fourcroydes*, and agave hybrid 11648 (*A. amaniensis* x *A. angustifolia*) under prolonged drought in field conditions. Leaf, stem, and root tissues were sequenced, and gene expression profiles were correlated with biomass composition, enzymatic hydrolysis of cell wall carbohydrates, histochemical analysis, and non-structural carbohydrates content.
- Differences in biomass accessibility were attributed to either lignin content or lignin composition, possibly through modification of s/g ratio promoted by changes in Caffeic Acid 3-O-Methyltransferase (COMT) transcript abundance. Unlike most plants, the most highly expressed transcripts do not encode photosynthetic proteins, but rather involved in stress response. Although the three cultivars presented quantitative differences in global gene expression, they activated a highly overlapping set of genes. The main molecular strategies employed by agave to cope with high-temperature and drought seem to consist in overexpressing HSP and LEA, as well as promoting raffinose accumulation as an osmolyte.
- In conclusion, our data provide vital new genetic information for the study of Agave species and provide new insights into cell wall architecture, recalcitrance, and resistance to abiotic stresses for these species.

## 1. BACKGROUND

Agaves are evergreen xerophyte monocots native to the semiarid regions of North America that are cultivated worldwide as ornamental plants (Nobel, 1994), and as crops for natural fiber and alcoholic beverages production (Davis & Long, 2015; Medina, 1954; Nobel, 1994). Among the agaves grown to obtain fiber, the most widely cultivated is *Agave sisalana* (Sisal), though other taxons such as *Agave fourcroydes* (henequen) and hybrid 11648 (*A. amaniensis* x *A. angustifolia*) are also of great importance (Davis & Long, 2015; Duarte et al., 2018; Medina, 1954). Brazil is currently the world’s largest producer and exporter of sisal fiber, representing 70% of the exports and 58% of the global production (Davis & Long, 2015; FAO, 2020). The Brazilian semiarid total area is around 83 million hectares (Projeto MapBiomas, 2019), which is bigger than most European countries, like France or Spain. Only a small portion of this territory is used for agriculture, and, in these areas, sisal is often the only growing crop alternative with attractive economic results (Alvarenga Jr., 2012; Broeren et al., 2017; Silva & Beltrão, 1999). Since the late 1960s, sisal fiber production has been decreasing dramatically, mainly due to competition with synthetic products. Nonetheless, in recent years, an emerging interest in natural fibers has made sisal production rise again (Davis & Long, 2015; FAO, 2020). This interest could be attributed to environmental concerns about microplastic pollution as well as the demand for niche markets (Cesa, Turra, & Baruque-Ramos, 2017; Davis & Long, 2015). Still, only 4% of the harvested leaves are converted into commercial fiber, resulting in a huge amount of bagasse that is dumped back into the field (Leão et al., 2016). This waste is rich in carbohydrates and organic acids, presents high technological potential (as pharmaceuticals, cosmetics and nanocellulose) and could be suited as biorefineries feedstocks (Borland, Griffiths, Hartwell, & Smith, 2009; Branco et al., 2010; Davis, Kuzmick, Niechayev, & Hunsaker, 2017; Dellaert, 2014; Lacerda, de Paula, Zambon, & Frollini, 2012; Michel-Cuello, Juárez-Flores, Aguirre-Rivera, & Pinos-Rodríguez, 2008; Morán, Alvarez, Cyras, & Vázquez, 2008; Santos, Vieira, Braz-Filho, & Branco, 2015).

The great success of Agave species in hostile environments is largely associated with its photosynthesis, the crassulacean acid metabolism (CAM) (Borland et al., 2009; Yang et al., 2015). CAM is a temporally regulated inorganic carbon concentration mechanism that relies on the primary CO2 fixation during nighttime, when evapotranspiration rates are reduced, to maximize water use efficiency, which results in up to 80% less water use when compared to conventional C3 and C4 crops (Borland et al., 2014; Stewart, 2015; Yin et al., 2018). Within CAM species, agaves are among the few that can achieve high productivity and, for that, are being considered as bioenergy feedstocks for dryland areas, which are commonly neglected and underused (Borland et al., 2014; Davis et al., 2017; Nobel, 1994; Owen, Fahy, & Griffiths, 2016; Somerville, Youngs, Taylor, Davis, & Long, 2010). Depending on region and species, commercially grown agave can achieve yields ranging from 8.5 to 22 Mg ha_−1_ (Davis, LeBauer, & Long, 2014); a recent field trial in Arizona (US) found yields between 4.0–9.3 Mg ha_−1_ yr_−1_ total biomass with 300–530 mm yr_−1_ of water input, which were already greater than conventional crops in Arizona, like cotton (1.5 Mg ha_−1_ yr_−1_), with less water consumption (Davis et al., 2017). However, theoretical analysis indicates even greater potential productivity (38 Mg (dry) ha_−1_ yr_−1_) (Owen et al., 2016). Also, agaves offer other desirable traits as biorenewables and biofuels feedstocks, such as the abundance of non-structural carbohydrates (mainly fructans), high shoot to root ratio, and low lignin content (Borland et al., 2009; Smith, 2008). Among cell wall polymers, lignin is the main cause of recalcitrance and a barrier for lignocellulosic biofuels production (Ragauskas et al., 2014; Simmons, Loqué, & Ralph, 2010). Therefore, understanding how the agave cell wall is formed may be important to find new ways to reduce lignin content without impairing plant growth (Simpson et al., 2011; Stewart, 2015; Vanholme, Ralph, et al., 2010). Furthermore, agaves constitute interesting models for the study of severe abiotic stress responses in plants, including drought and high temperatures, which is relevant for the development of agronomic solutions associated with climate change and bioenergy production (Borland et al., 2009; Stewart, 2015).

Despite their social and economic importance, few studies have been carried out with *Agave* species and their hybrids, especially at the molecular level. This is mainly because of: (i) lack of basic genetic knowledge; (ii) large genomes, which are estimated to be between 2,940 and 4,704 Mbp; (iii) long life cycles (5-12 years) (Davis & Long, 2015; Simpson et al., 2011). Nonetheless, the *Agave* genus is rich in genetic diversity, as ploidy varies from 2*n* to 8*n* (*n*=30) even within species, and hybridization has occurred frequently in wild cultivars, contributing to increasing variation in ploidy and complexity of hybrid genomes (Davis & Long, 2015; Simpson et al., 2011). Although DNA-seq data from *Agave tequilana* has been deposited at NCBI, a reference genome sequence has not been published to date. Furthermore, only six studies have been published on agave transcriptomics, but none of them has explored the molecular basis of cell wall biosynthesis (Abraham et al., 2016; Cervantes-Pérez et al., 2018; Gross et al., 2013; Huang et al., 2018, 2019; McKain et al., 2012; Sarwar et al., 2019; Simpson et al., 2011).

In the present work, we analyze the transcriptomic profile of *Agave sisalana*, *Agave fourcroydes*, and agave hybrid 11648 (*A. amaniensis* x *A. angustifolia*) in field conditions under prolonged drought, and correlate transcription with non-structural carbohydrates profiles, biomass composition, enzymatic hydrolysis of cell wall carbohydrates, histochemical analysis of leaf anatomy. The valuable resources generated offer a new window of opportunity to molecular breeders fulfill the emerging expectations for agave use as biorefineries feedstocks for dryland areas and provide insights into abiotic stress resistance and cell wall architecture.

## 2. METHODS

### 2.1. Plant material and growth conditions

*A. sisalana*, *A. fourcroydes*, and agave hybrid 11648 (*A. amaniensis* x *A. angustifolia*) (H11648) samples were harvested under field conditions from Embrapa’s germplasm bank, located on the Experimental Station of Monteiro-PB, Brazil (7° 53’ south latitude, 37° 07’ west longitude, 619 m altitude). Plants were grown without artificial irrigation in non-calcic brown soil and semi-arid climate (BS, according to the Köppen system) (BRASIL, 1972; KÖPPEN & GEIGER, 1936). Rainfall data from the National Institute of Meteorology of Brazil shows that the municipality of Monteiro has suffered from a prolonged drought, not exceeding 150 mm of rainfall, since April 2014 until the sampling date (supplementary material S1). For all cultivars, leaf (central fraction), root, and stem were sampled from seven-year-old healthy adult plants growing side-by-side (n=3). As agave leaves grow during daytime (Abraham et al., 2016), sample harvested at mid-day were selected for sequencing to better dissect the relationship between abiotic stress response and cell wall biogenesis.

### 2.2. RNA Extraction and sequencing

Total RNA was extracted according to the protocol described by Zeng & Yang, 2002, with the modifications proposed by Le Provost *et al.*, 2003. The RNA concentration and quality were verified using a Nanodrop 2000 spectrophotometer (Thermo Scientific) and integrity with HT RNA LabChip® Kit (Caliper Life Sciences). mRNA libraries and sequencing were done at the High-Throughput Sequencing Facility of the Carolina Center for Genome Sciences (University of North Carolina at Chapel Hill, USA). The libraries were prepared using the KAPA Stranded mRNA-Seq kit (07962193001) for Illumina platforms following the manufacturer’s protocol, using 1 μg total RNA. To facilitate the assembly of transcripts, larger insert sizes (150-180 bp) were generated altering the fragmentation step of the protocol to 7 min at 94°C. The sequencing was done on the Illumina HiSeq 4000 system. Libraries were separated into two pools and each pool was sequenced in a single lane, generating 50bp paired-end reads.

### 2.3. *De novo* transcriptome assembly, quantification, and annotation

FastQC software was used for quality control (Andrews, 2010). Reads from each cultivar were assembled separately using Trinity v. 2.5.1 (Grabherr et al., 2011). Transcript quantification was performed using kallisto v 0.44.0 (Bray, Pimentel, Melsted, & Pachter, 2016) with 100 bootstraps, returning TPM (Transcripts Per Million) values representing the transcripts’ abundance. For ORF prediction, we used Transdecoder v. 5.0.2 (Haas et al., 2013) configured to a minimum length of 200 nucleotides. Subsequently, we selected the longest isoform of each locus, considering only those with a TPM value greater than 1, and ORF length longer than 255 nucleotides. Functional annotation assignment was performed using Pannzer2 (Törönen, Medlar, & Holm, 2018). Unannotated proteins were blasted (BLASTp) against the Uniref90 database.

### 2.4. Fungal identification and classification

We have developed an in-house Perl script that used the Uniref90 BLASTp results and a list of fungal genera from Taxonomy DB (NCBI) to separate the fungal transcripts from the agave ones. The criterion to classify as fungal sequence was based on the percentage of fungal genera in the top blast hits considering an e-value <= 1e-10 (it classifies as fungi if at least 80% of the top ten hits are fungi).

To confirm this classification and separate the fungi transcripts between Ascomycota, Basidiomycota and other fungi, we used two strategies: a) Kaiju program v. 1.6.3 (Menzel, Ng, & Krogh, 2016) with fungi database to perform the taxonomy composition analysis and b) BLASTn against three separate databases of coding sequences (CDS) obtained from NCBI (Ascomycota genomes, Basidiomycota genomes and complete genomes of other fungal phyla) that were analyzed by an in-house Perl script, considering the alignment query coverage >= 30% and the e-value <= 1e-20 to separate the fungi groups.

### 2.5. Tissue-specific transcripts analysis

To analyze the specific transcription profile of each tissue, we used the SPM (specificity measure) metric implemented by the software tspex (https://github.com/apcamargo/tspex). Tissue specific genes with SPM >= 0,95 were selected.

### 2.6. Orthologous analysis between cultivars

To compare the transcriptome within cultivars, we first defined orthologous groups using OrthoMCL v. 1.4 (L. Li, 2003) using nucleotide sequences from the assembled transcriptomes.

### 2.7. Differential gene expression analysis and GO enrichment

R package sleuth v. 0.29.0 (Pimentel, Bray, Puente, Melsted, & Pachter, 2017) was used for the differential expression analysis with the Wald test. Pairwise comparisons were made between tissues within each cultivar, and differentially expressed genes (DEG) with a fold change > 2 and FDR ≤ 0.05 were selected. Representative DEG from each tissue (using sets with two lists of genes preferentially expressed in that tissue) were submmited to Gene Ontology enrichment analysis using topGO R package (Alexa, Rahnenfuhrer, & Lengauer, 2006). We considered only enriched Gene Ontology terms with p-value < 0.05 for further analyses.

### 2.8. Comparative genomics

To identify exclusive and expanded gene families from *Agave* and other related species, we employed a comparative genomics approach using OrthoFinder v. 2.3.1 (Emms & Kelly, 2015). Single-copy orthologs were aligned with MAFFT v. 7.394 (Katoh & Standley, 2013) configured to use the L-INS algorithm with 1,000 iterations. All alignments were concatenated in a supermatrix that was used for the phylogenetic inference with IQ-TREE v. 1.6.8 (Nguyen, Schmidt, von Haeseler, & Minh, 2015). BadiRate v. 1.35 (Librado, Vieira, & Rozas, 2012) was used to identify expanded gene families using a FDR threshold of 0.05. We compared our data to protein sequences from genomes available at the Phytozome database (Goodstein et al., 2012). We selected *Amborella trichopoda* (outgroup) v. 1.0, *Asparagus officinalis* v. 1.1, *Eucalyptus grandis* v. 2.0, *Sorghum bicolor* v. 3.1.1, *Ananas comosus* v. 3, *Punica granatum* (Qin et al., 2017) and *Saccharum spontaneum* (Zhang et al., 2018). These species were chosen based on their photosynthetic metabolism, biomass composition and productivity.

### 2.9. Non-structural carbohydrates profiling

Rhamnose, ribose, fructose, pinitol, galactose, mannitol, sorbitol, glucose, sucrose, maltose, trehalose, melibiose, and raffinose were quantified in leaves and stem using a gas chromatographer (Hewlett-Packard® 6890) connected to a quadrupole mass spectrometer (Hewlett-Packard® model 5973) following the same methodology as Freschi *et al.*, 2010. The initial running condition was 95 °C for 2 min, followed by a gradient up to 320 °C at 8 °C min_−1_. The column used for separation was an HP-1701 (30 m, I.D. 0.25 mm, 0.5 μm), with helium as the carrier gas, with a flux of 4 mL min_−1_. The endogenous metabolite concentration was obtained by comparing the peak areas of the chromatograms with commercial standards. All measurements were made in triplicate.

Water extraction for 15 min at 80°C (Mancilla-Margalli & López, 2006) followed by thermally hydrolyzation (80°C for 30h in water) of half of the samples (Michel-Cuello et al., 2008) was performed to estimate of the oligosaccharides and monosaccharides within the cultivars. Oligosaccharides were estimated by comparing the difference between the hydrolyzed and non-hydrolyzed fractions. All the samples were evaluated using reducing sugars content measured with DNS assay (Miller, 1959).

### 2.10. Determination of structural carbohydrates and lignin in biomass

Cellulose, hemicellulose, lignin, extractives and ash were quantified according to the standardized analytical methods of Sluiter *et al.*, 2016.

### 2.11. Enzymatic hydrolysis of cell wall carbohydrates

A simple enzymatic hydrolysis was performed to verify the saccharification and accessibility potential of the agave biomass. We performed the enzymatic hydrolysis assays following the Bragatto *et al.*, 2012 protocol with the modifications of Lepikson-Neto *et al.*, 2014. Cellulase enzyme C2730 (Sigma) was used, and the content of reducing sugars was measure with DNS assay (Miller, 1959), as suggested by Lepikson-Neto *et al.*, 2014. Also, for comparison, we added an Energy Cane sample (US59-6; whole plant biomass) processed in parallel.

### 2.12. Histochemical analysis of Agave leaves

To complement the compositional data, the central fraction of fully developed leaves from one-year-old plants of the three Agave cultivars was sampled to verify the distribution of lignin, pectic compounds, and callose. Leaves cross-sections were stained with phloroglucinol-HCl, for lignin detection (Johansen, 1940), and with ruthenium red, for pectic compounds (Johansen, 1940), and, then, observed under light Olympus BX 51 photomicroscope equipped with an Olympus DP71 camera. For callose, samples were stained with aniline blue (Currier & Strugger, 1956) and observed under an epifluorescence microscope with a UV filter (BP 340 to 380 nm, LP 425 nm).

## 3. RESULTS

### 3.1. Transcriptome assembly and annotation

The total number of transcripts obtained by *de novo* assembly of RNA-seq reads from the different *Agave* species ranged from 136,692 to 170,474. After the removal of short (<250 bp) and weakly expressed transcripts (TPM<1), we selected one transcript per locus. The total number of transcripts ranged from 23,973 to 26,842. These values are similar to those reported by Gross *et al.*, 2013 in the transcriptomic analysis of other agave species: 34,870 and 35,086 protein-coding loci for *A. tequilana* and *A. deserti*, respectively. In addition, *Asparagus officinalis,* which is the closest specie to the agaves with a reference genome available, has 27,656 protein-coding genes (Harkess et al., 2017). Several unannotated proteins ranging from 12,4 to 14,2% were found. All transcriptomic data is available in supplementary table 2 (S2). Remarkably, around 12% of the root transcripts belonged to fungi. The presence of these fungal transcripts was persistent in roots for all cultivars and biological replicates, and 99.8% of those transcripts were root-specific, with SPM > 0.95. Interestingly, 5-6% of the fungal transcriptomes were annotated as heat shock proteins - HSP, and, in some cases, the fungal HSP were as abundant as some of the agave’s HSP (depending on the cultivar, the most expressed fungal HSP ranged from 120 to 413 TPM.). It was possible to identify 46.8-54.7% of the fungal transcripts as Ascomycetes and 16.3-20.6% as Basidiomycetes. The fungal transcripts were excluded from further analysis.

#### 3.1.1. Most expressed transcripts

From the transcriptome quantification, we have ranked the ten most highly expressed transcripts in each tissue for each cultivar (Table 1). Surprisingly, a similar set of highly expressed transcripts was identified in all tissues of the three cultivars. One of those transcripts is *LEA-5* (late embryogenesis abundant protein 5). *LEA* is one of the most expressed, with transcription ranging from 2,772 to 15,947. *LEA* transcripts have been previously found in many vegetative tissues with several functions associated with abiotic stress responses, including heat and salinity in *Agave* species (Liang et al., 2014; Pedrosa, Martins, Gonçalves, & Costa, 2015; Tamayo-Ordóñez et al., 2016). Other highly expressed transcripts encode Ubiquitin and Polyubiquitin proteins. However, for H1648 leaves and stem, these transcripts were not at the topmost. Ubiquitins are related to proteolysis signaling, which has also been associated with drought stress (Flick & Kaiser, 2012a; Lyzenga & Stone, 2012). Many heat shock proteins and uncharacterized protein transcripts are highly expressed in the three cultivars as well.

**Table 1.**
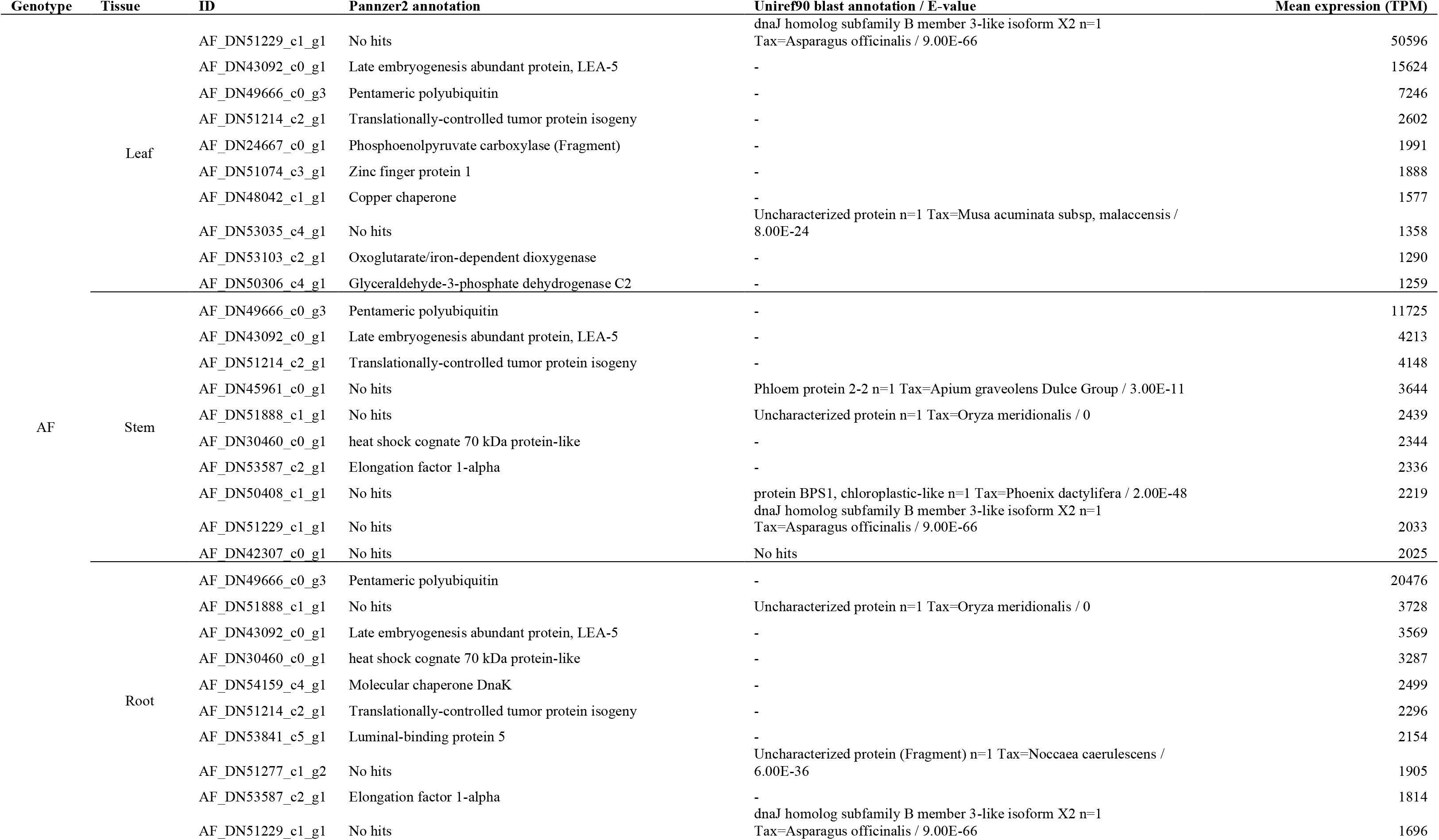
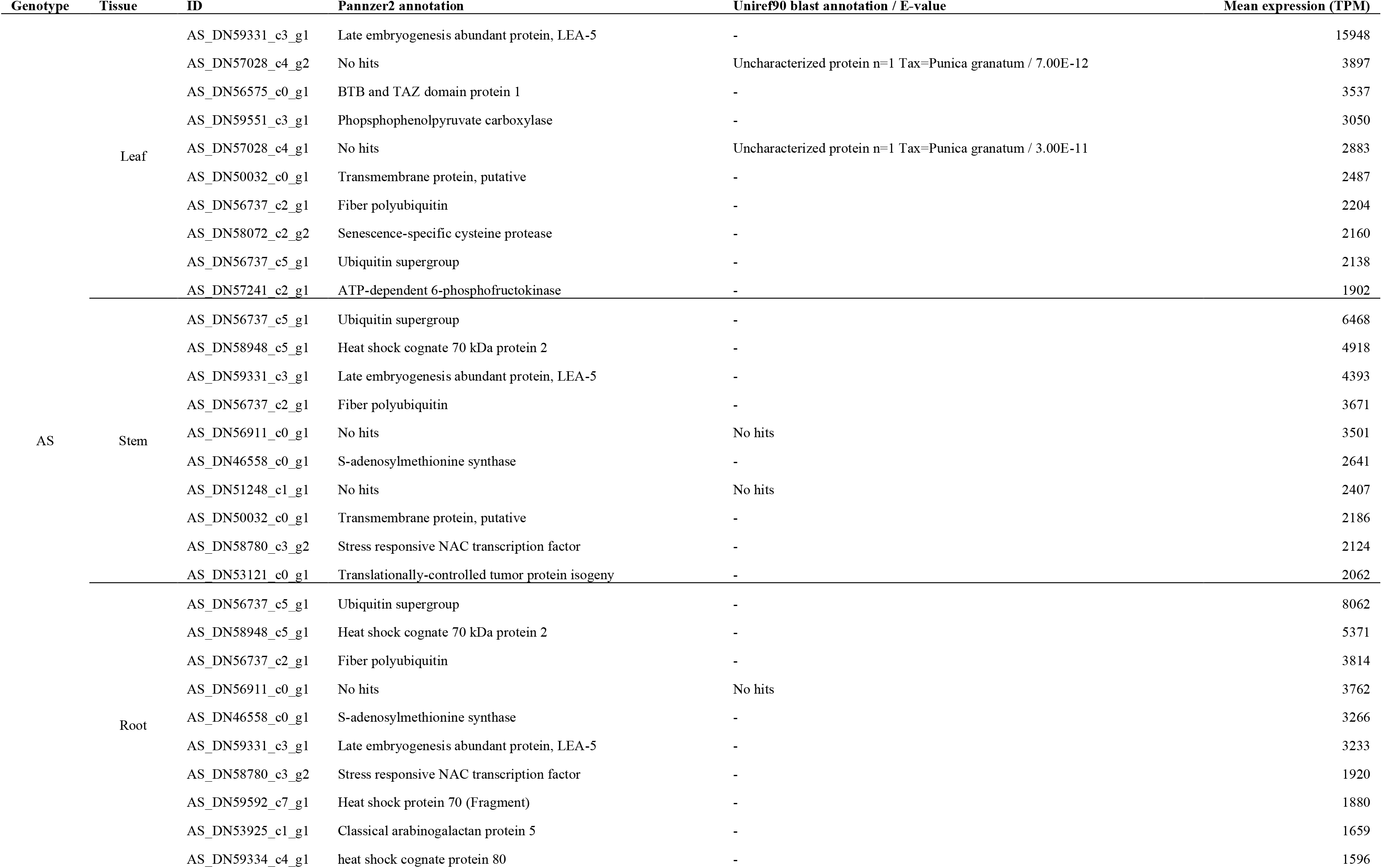
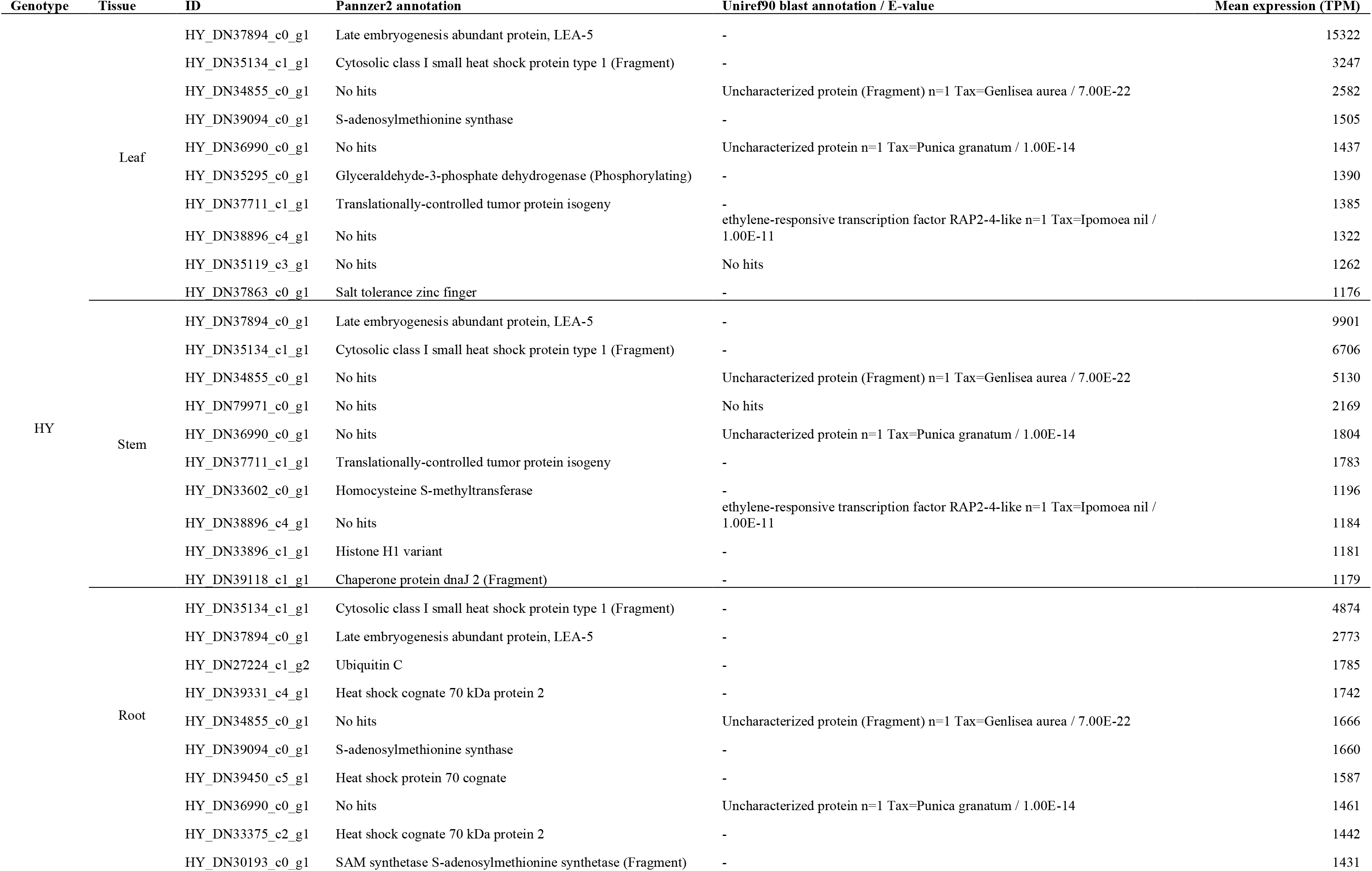
The most highly expressed transcripts in each tissue for the three cultivars. The annotation was performed by Pannzer software and BLASTp against Uniref90. Uniref90’s annotations are shown only for Pannzer’s no hits.

#### 3.1.2. Tissue-specific transcripts

As expected, our analysis revealed that most of the leaf-specific transcripts are photosynthesis-related (Table 2). Interestingly, in the stem for all cultivars, we have found several homeobox transcripts, which are proteins commonly associated with development (Mukherjee, Brocchieri, & Burglin, 2009). The most expressed root-specific genes are no-hit proteins, and for *A. fourcroydes* and *A. sisalana* ubiquitins are present as well. Also, *A. sisalana* has one root-specific heat shock protein with high expression; and H11648 has a Thaumatin, which is a sweet-tasting protein homologous to pathogenesis-related (PR) protein PR-5 (Min et al., 2003). One member of the PR-5 protein family, Osmotin, is accumulated in cells adapted to osmotic stress (Singh et al., 1987).

**Table 2.**
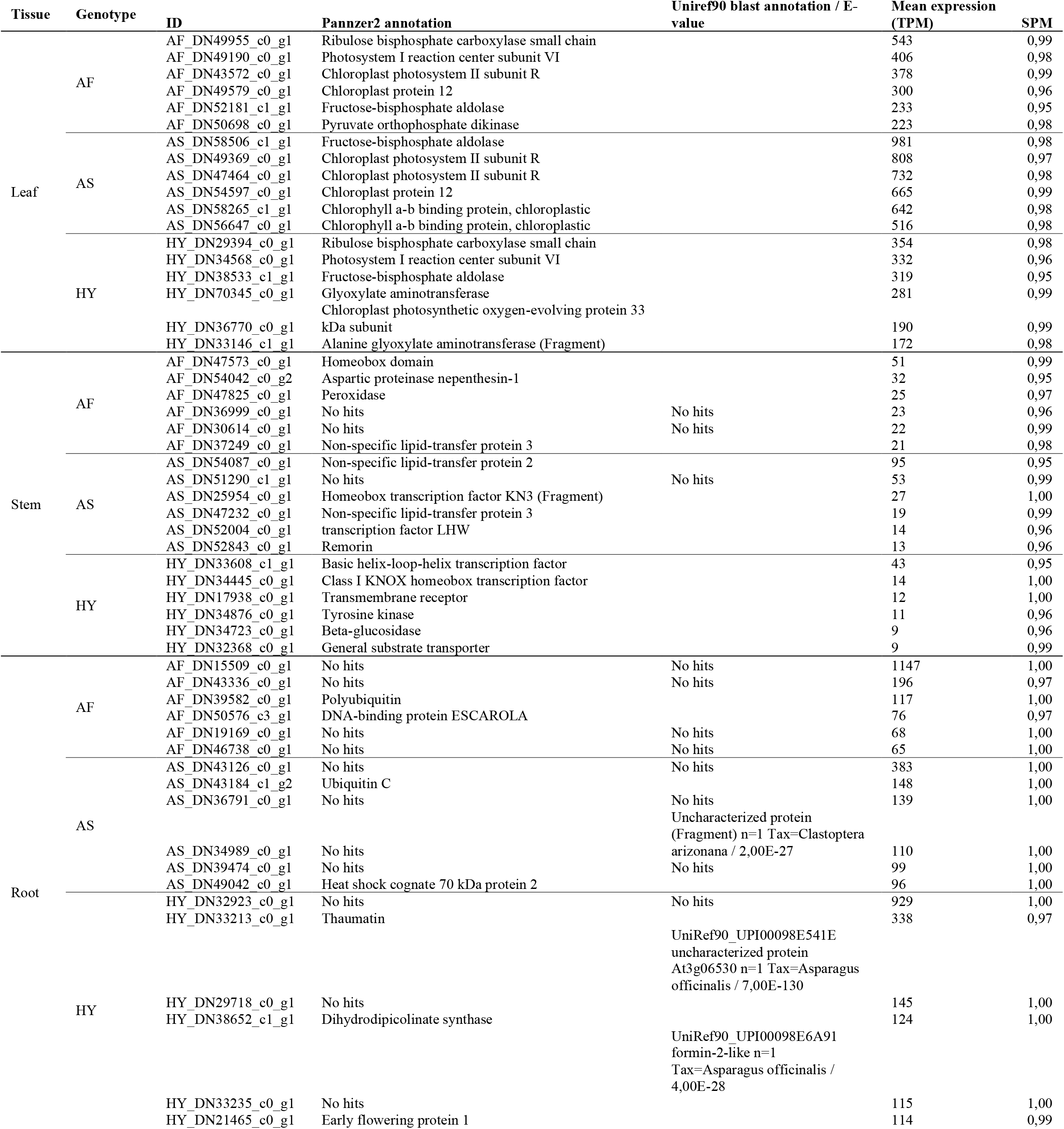
Six most expressed tissue specific transcripts for the three cultivars. The annotation was performed by Pannzer software and BLASTp against Uniref90. Uniref90’s annotations are shown only for Pannzer’s no hits.

#### 3.1.3. Phenylpropanoids pathway

The phenylpropanoid pathway is composed of many branches, but all of them share the same common precursors (Fraser & Chapple, 2011). We analyzed the transcriptional profile of the lignin and flavonoids biosynthesis branches (Figure 6a). At least one member of each gene family of the phenylpropanoid pathway was differentially expressed, and here we highlight the main focal points. Phenylalanine Ammonium Lyase (PAL), the first enzyme of the pathway, presented similar behavior in *A. fourcroydes* and *A. sisalana*, in both species, the most expressed isozyme gene had higher transcription in the stem. Also, an orphan PAL isozyme gene was found for each species. Notwithstanding, for H11648, the main PAL isozyme gene was not differentially expressed. The first diverging point is the conversion of p-coumaroyl CoA by either Chalcone Synthase (CHS) or Hydroxycinnamoyl-coenzyme A Shikimate:Quinate Hydroxycinnamoyltransferase (HCT). For CHS, which is the starting enzyme for the flavonoid biosynthesis, two main encoding genes were found to be differentially expressed. However, the overall expression of these two isozymes was relatively low. Curiously, one of those isozymes presented significantly higher expression at the H11648 stem, although, its transcription was not very high. For HCT, which catalyzes the outset of the lignin branch, four HCT isozyme genes were deferentially expressed, and two were responsible for the majority of the transcription. Whereas for H11648, these two isozyme-enconding gene did not present any relevant differences. For the other cultivars, each HCT isozyme-encoding transcript behaved oppositely, being either more expressed in leaves or stems; a phenomenon particularly evident in *A. sisalana*. On the phenylpropanoid pathway, there are only two enzymes exclusive to syringyl (S) monomer biosynthesis: Ferulate 5-Hydroxylase (F5H) and Caffeic Acid 3-O-Methyltransferase (COMT). We identified just one F5H isozyme-encoding transcript differentially expressed and more expressed in stems than leaves in all cultivars. On the other hand, three COMT isozymes were found, with two predominantly expressed in stems and third one mainly found in leaves. Interestingly, in general, COMT expression was higher at *A. fourcroydes* stem.

**Figure 1.**
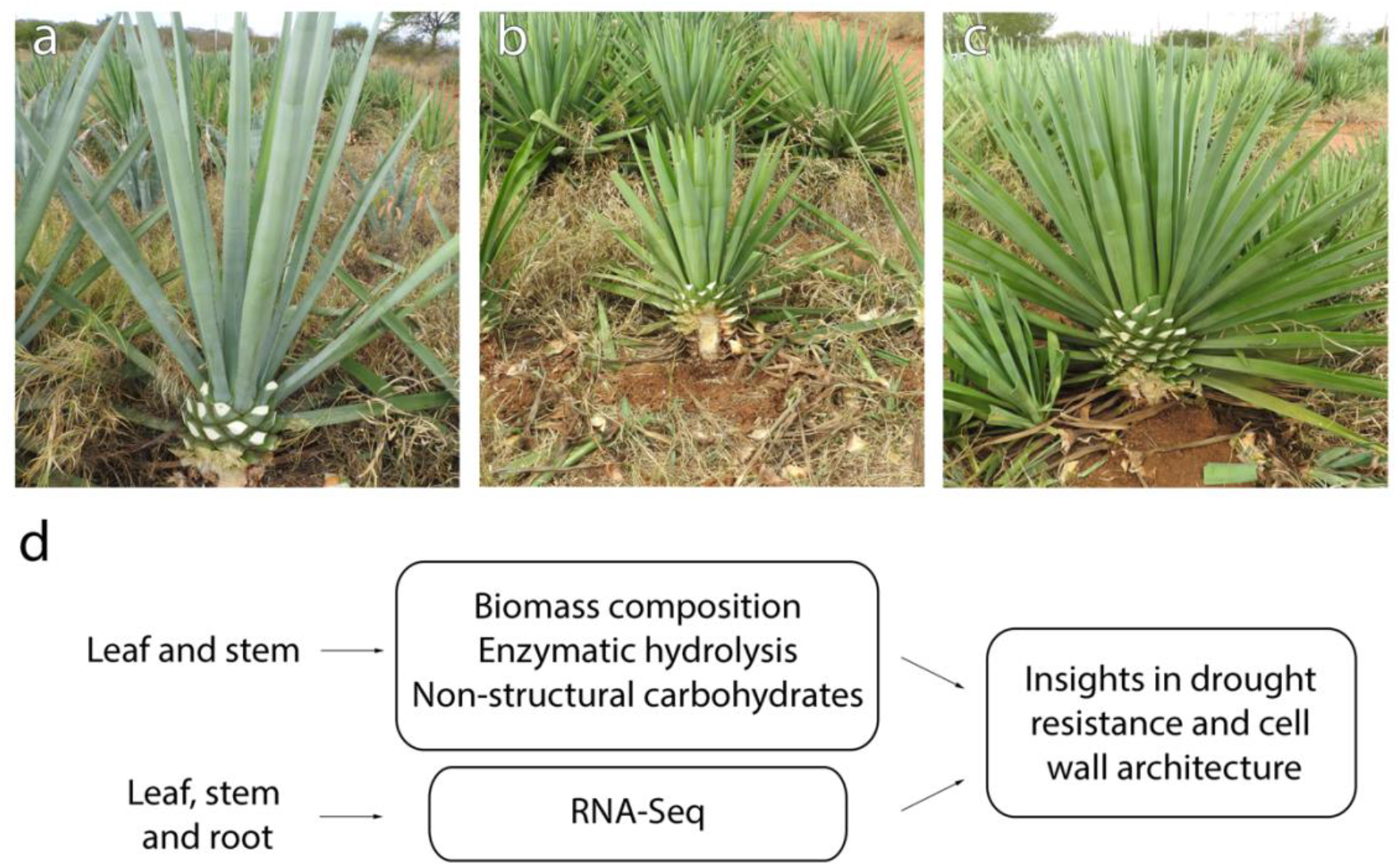
The three cultivars sampled at Embrapa’s germplasm bank and a summary of the methods used in this work. (a) *A. fourcroydes*; (b) *A. sisalana*; and (c) H11648. Using samples of three different individuals of these three cultivars, we correlated chemical and transcriptomic data to provide new insights into cell wall architecture, recalcitrance, and resistance to abiotic stresses.

**Figure 2.**
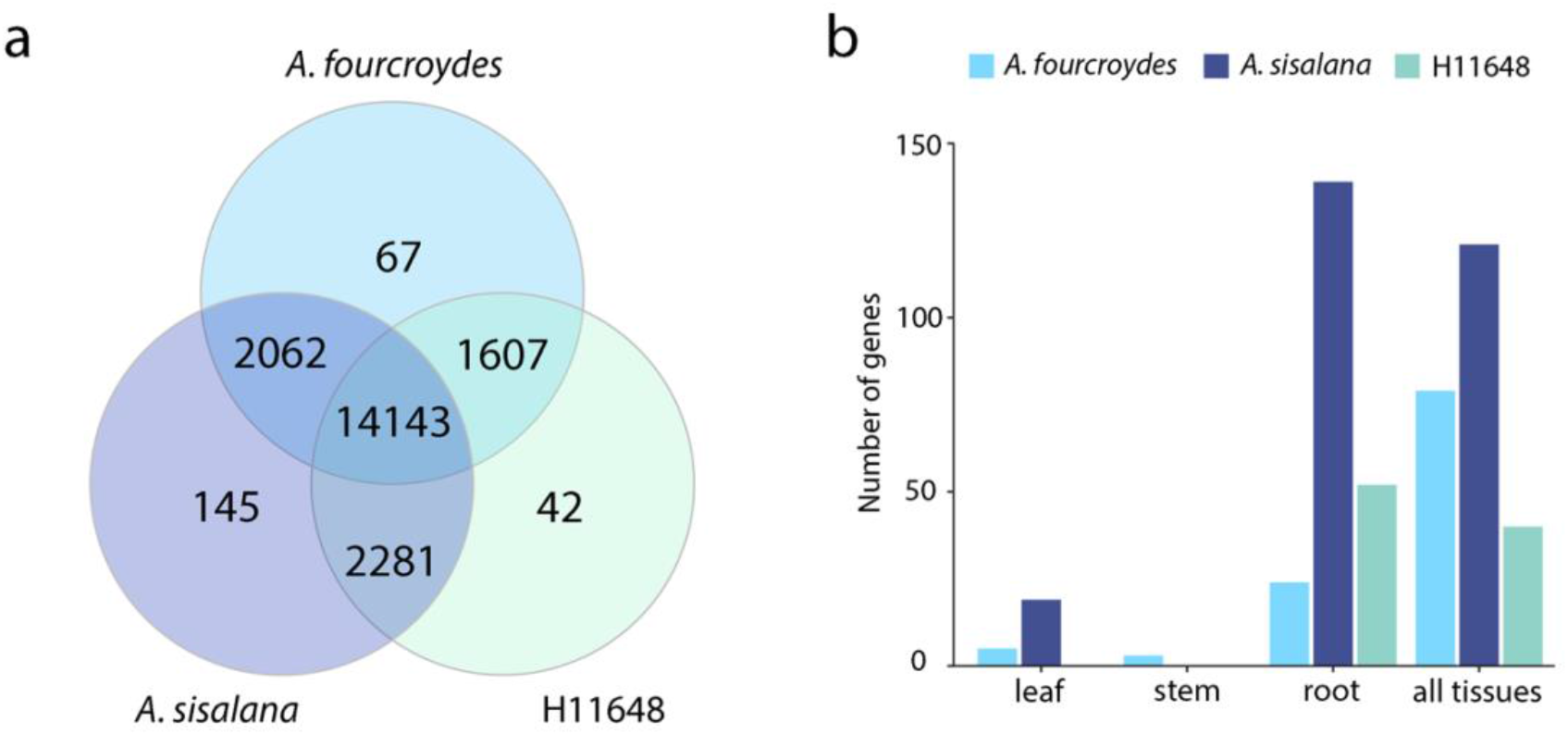
(a) Venn diagram showing number of shared and exclusive gene families (nucleotides) between the assemblies of the three cultivars. (b) Number of exclusive genes per tissue and cultivars.

**Figure 3.**
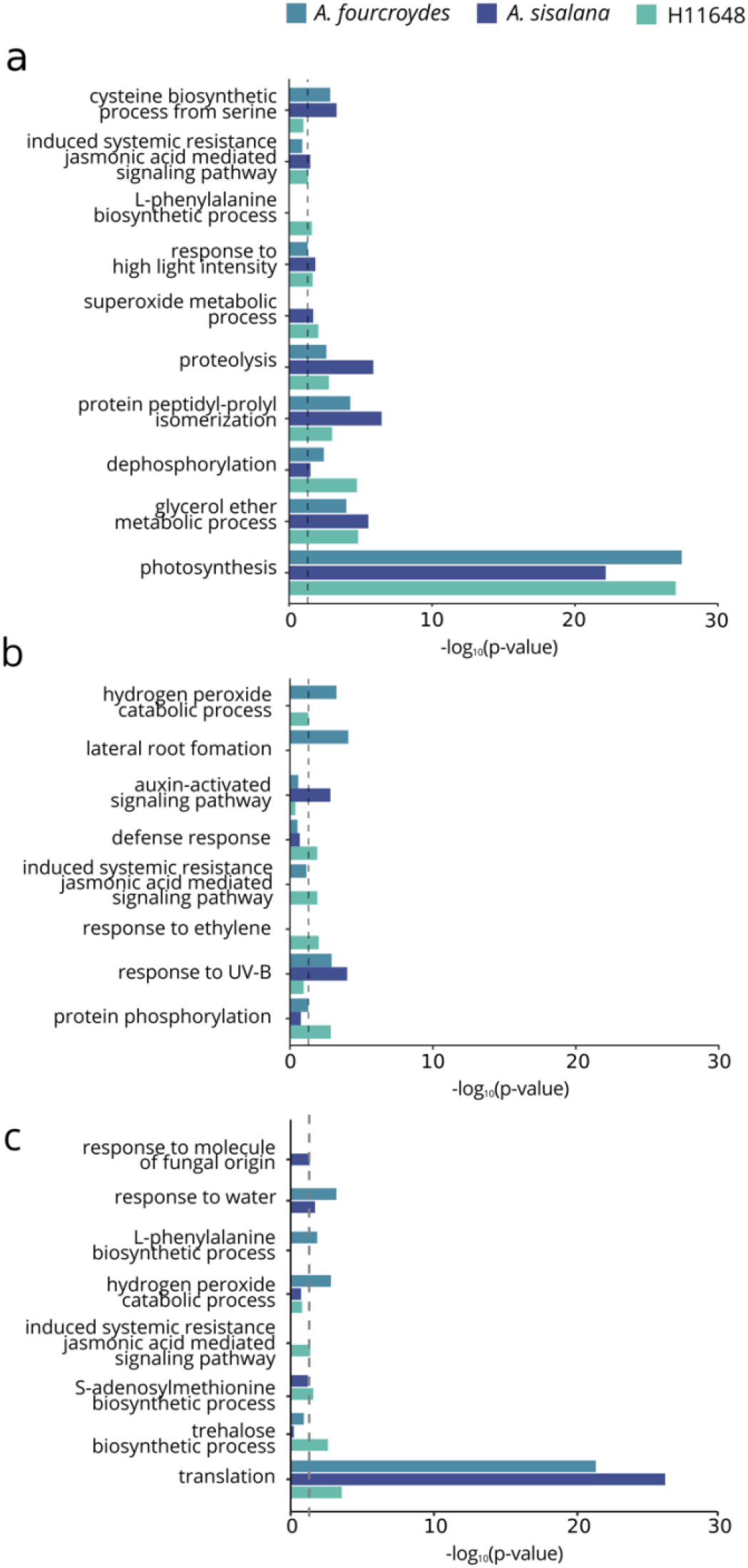
Significant (p-value < 0,05) Gene Ontology terms enriched in leaf (a), stem (b) and root (c) for the three cultivars. The dashed line shows the significance threshold adopted.

**Figure 4.**
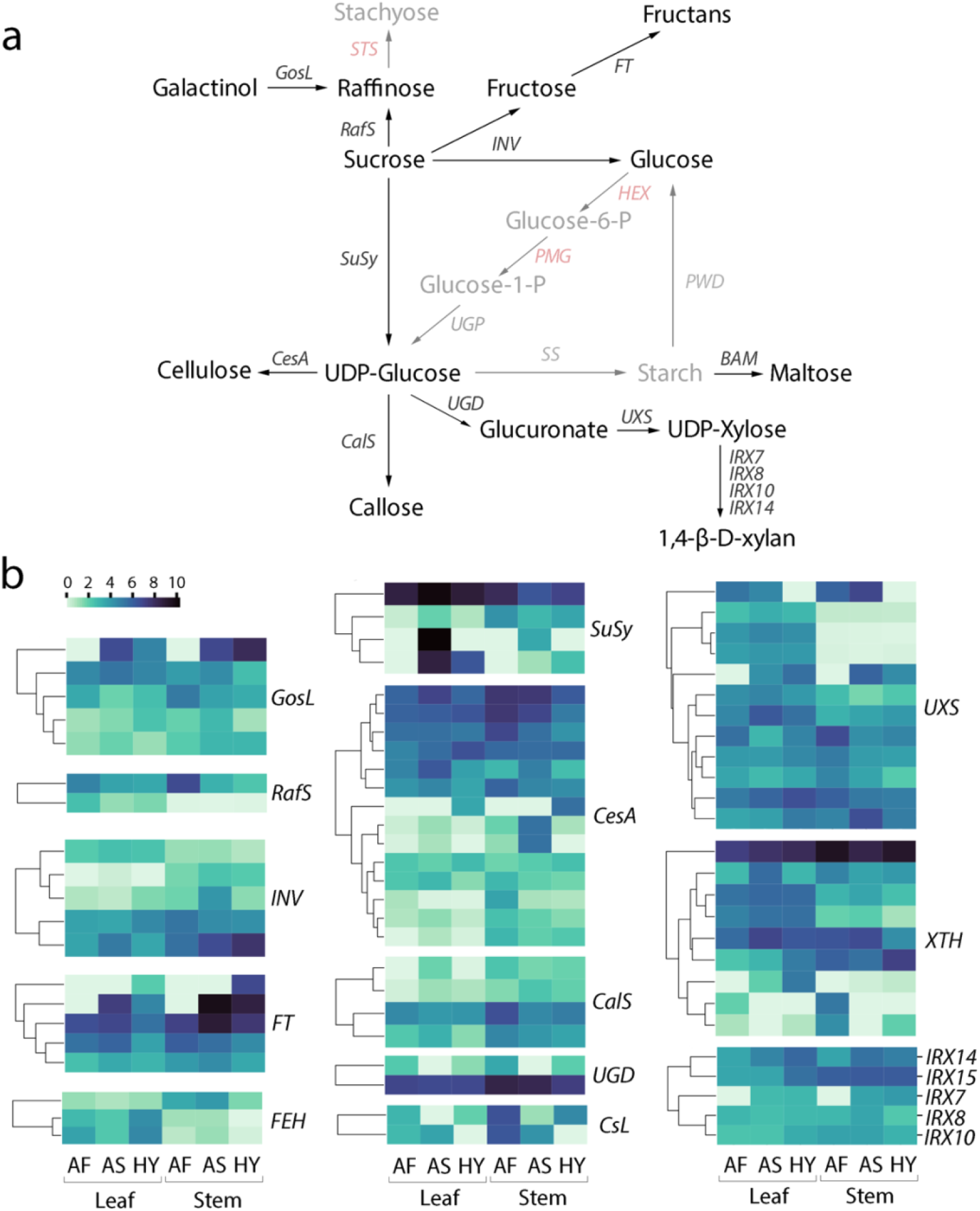
Gene expression of cellulose and associated carbohydrates pathways. (a) Schematic representation of the pathways, showing the coding genes for each enzyme. Red and grey indicate genes absent in our dataset and expressed at very low levels, respectively. (b) Heatmaps of log2 normalized values, representing the gene expression value in each tissue. AF: *A. fourcroydes*, AS: *A. sisalana*, HY: H11648.

**Figure 5.**
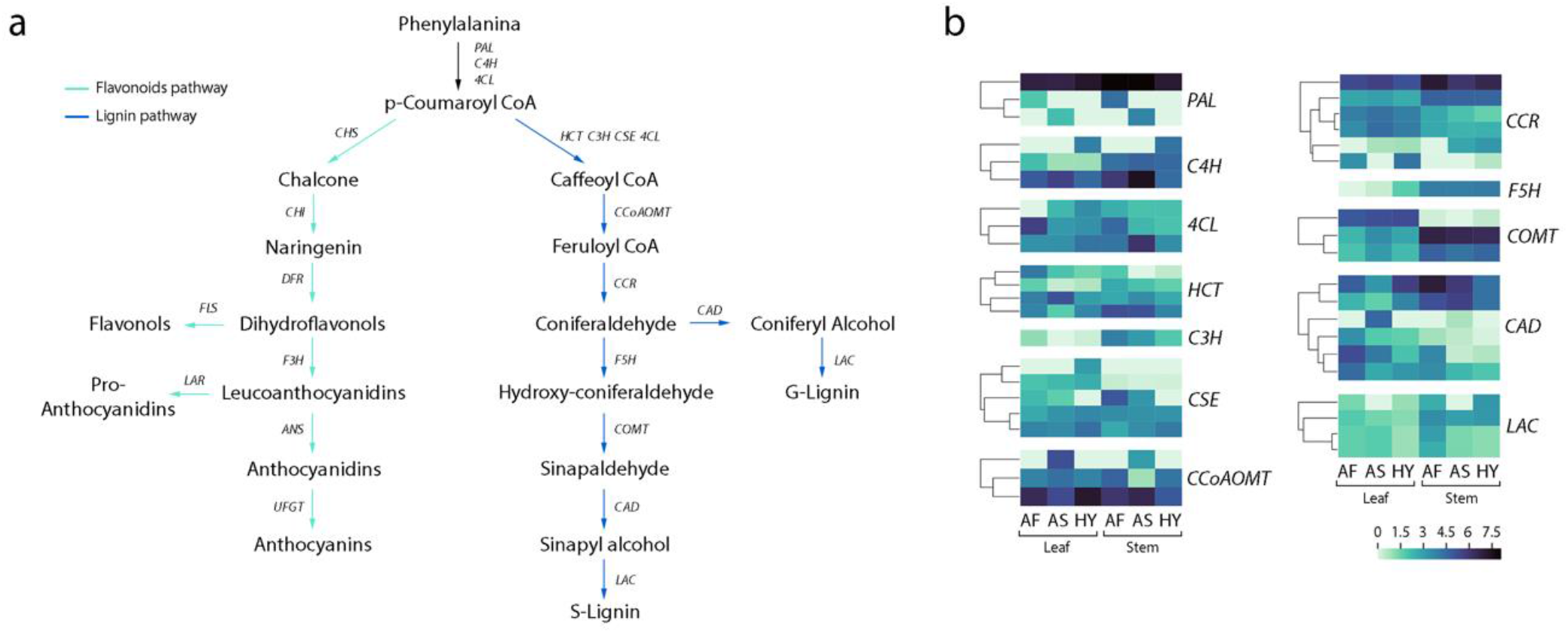
Gene expression of the phenylpropanoids pathway. (a) illustrates the pathway, showing the flavonoids (green arrows) and lignin (blue arrows) branches. (b) Heatmaps of log2 normalized values, representing gene expression value in each tissue for the lignin pathway genes. Flavonoids pathway genes did not present a very high expression and are not shown here. AF: *A. fourcroydes*, AS: *A. sisalana*, HY: H11648.

**Figure 6:**
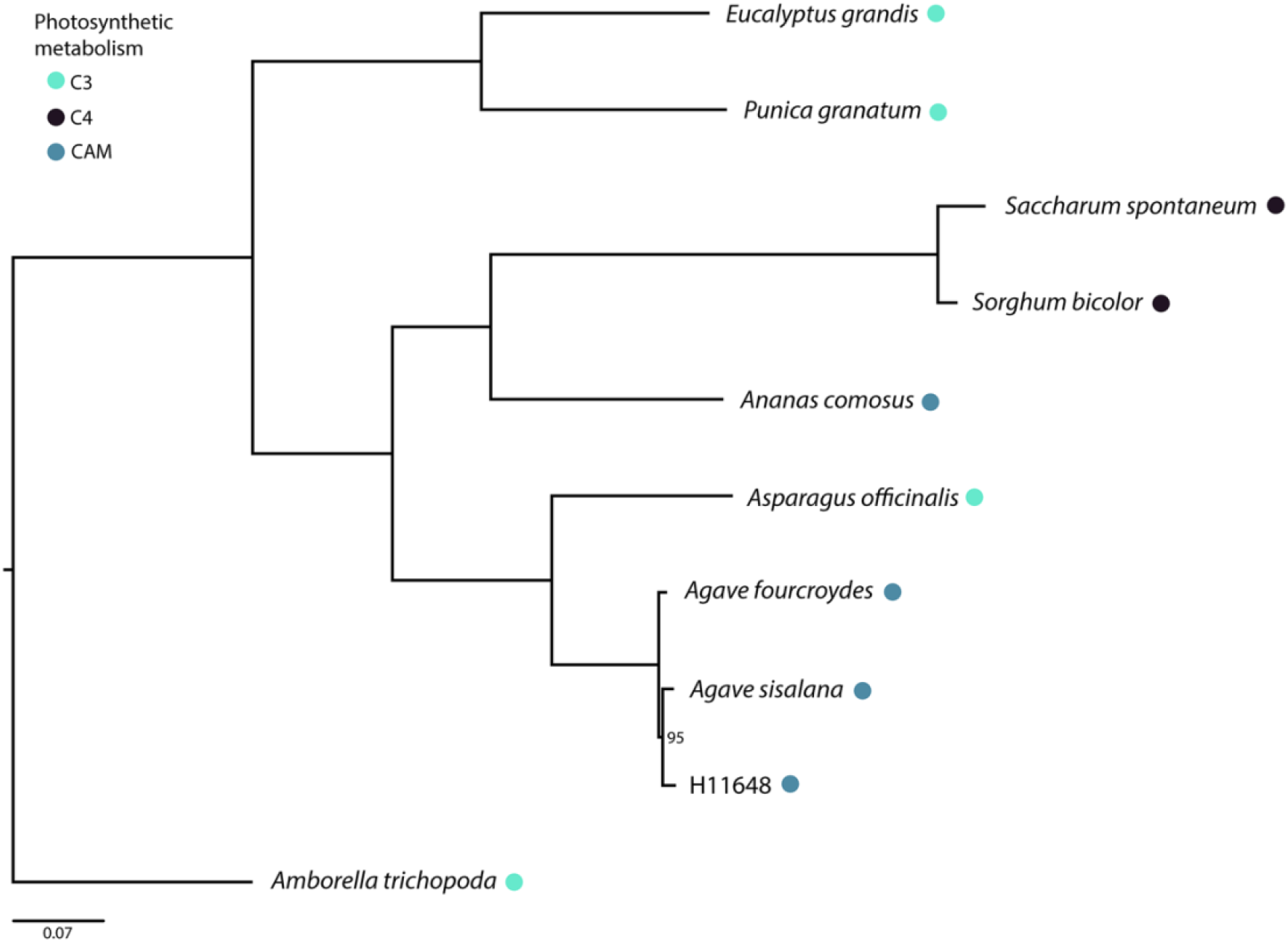
Maximum likelihood phylogenetic inference for 10 plant species. *Amborella trichopoda* was used as outgroup. Tree was reconstructed using 978 single-copy ortholog groups in IQ-TREE with 1,000 bootstrap randomizations. Phylogeny is scaled in substitutions per positions.

#### 3.1.4. Exclusive and expanded gene families of Agave

Twelve expanded families were found in the *Agave* clade using phylogeny and the birth and death models to calculate rates of gains and loss of genes through evolution (see methods 2.8). Among them, we identified a few transcription factors like MYB, Far-Red Impaired Response 1 (FAR1) and Zinc finger BED domain-containing protein DAYSLEEPER (OG0000003, OG0000022, and OG0000084, respectively). FAR1 activates the transcription of *Far-Red Elongated Hypocotyl 1* (*FHY1*) and its homolog, *FHY1-Like* (*FHL*), and positively regulates chlorophyll biosynthesis via the activation of *Delta-aminolevulinic acid dehydratase 1* (*HEMB1*) (Hudson, Lisch, & Quail, 2003; Lin et al., 2007; Tang et al., 2012; Wang & Deng, 2002). Two expanded families unannotated (OG0000281 and OG0000512) and one alpha expansin were also detected. No heat shock proteins or other apparent abiotic stress-related families were expanded.

Regarding biomass, we have found some important exclusive families. Gene families that contain at least one representative member shared in the agave cultivars and in the other analyzed species were considered as exclusive families. Among the 149 that are shared with *Saccharum spontaneum*, 68 are unannotated, and two are directly related to the cell wall: callose synthase 5 (OG0016451) and cellulose synthase (fragment) (OG0013945). In comparison, *Eucalyptus grandis* 94 exclusive families were found, of which only 48 were annotated. We identified two families related to lignin metabolism: a Caffeoyl-CoA 3-O-methyltransferase (CCoAMT) (OG0014109) and Laccase-14 (LAC14) (OG0013133). We also identified an endoglucanase (OG0012311) and an alpha-glucosidase (OG0012714). The full list of ortholog genes families generated by the comparative genomics approach is presented at the supplementary table 3 (S3).

### 3.2. Chemical analysis

#### 3.2.1. Non-structural carbohydrates profile

The GC-MS analysis detected all the following saccharides in both leaf and stem: fructose, glucose, sucrose and raffinose (Fig. 7a). Among them, raffinose was the most abundant carbohydrate for every cultivar and tissues sampled. Raffinose is an oligosaccharide related to osmotic and oxidative stress response in plants. Under dry conditions, raffinose acts as a compatible osmolyte avoiding water loss to the environment (Ende, 2013; Nishizawa, Yabuta, & Shigeoka, 2008; Sengupta, Mukherjee, Basak, & Majumder, 2015). In seeds of monocots, such as barley, corn, and sorghum, raffinose concentrations oscillates between 2.6-7.9 mg/g (Kuo, VanMiddlesworth, & Wolf, 1988). In contrast, for *Agave* vegetative tissues, we encountered mean values ranging from 10.96 to 23.38mg/g. Perhaps, the elevated presence of this carbohydrate could be related to the prolonged drought that challenged the plants under field conditions. Interestingly, no significant differences were found between the samples for this carbohydrate. That was not the case for fructose, glucose and sucrose. Glucose content was statistically different in the comparison between *A. fourcroydes* and H11648 leaves, with almost double of this carbohydrate being present in the leaves of H11648. However, there were no significant differences for the other carbohydrates nor between *A. sisalana* and *A. fourcroydes*. Among the leaves of *A. sisalana* and H11648, a difference was detected in glucose and fructose, again almost 50% more was quantified in the leaves of the hybrid.

**Figure 7.**
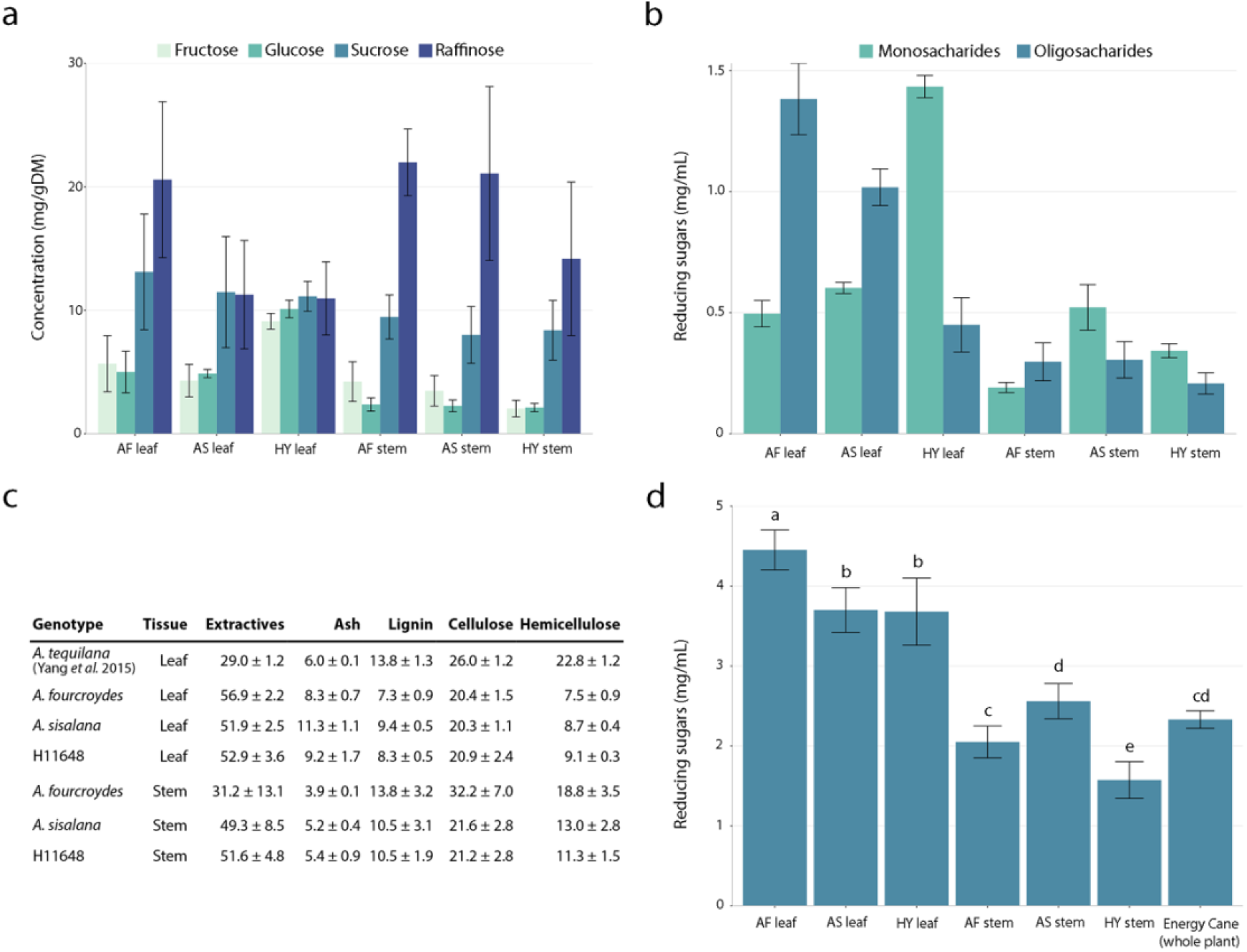
(a) Soluble carbohydrates profile obtained by GCMS; (b) Estimate of the oligosaccharides and monosaccharides fractions; (c) Compositional analysis of Agave leaves and stem, values are shown in percentage; (d) Enzymatic hydrolysis of cell wall carbohydrates. AF: *A. fourcroydes*, AS: *A. sisalana* and HY: H11648.

The highest concentrations of water-soluble carbohydrates were found in leaves regardless of the cultivar (Fig. 7b). *A. fourcroydes* and H11648 contained the highest sugar content. However, their mono and oligosaccharides fractions were contrasting. For *A. fourcroydes*, 26.41% of sugars detected were monosaccharides, while the same category in the H11648 represented 76.11%,. Interestingly, the transcriptomic data revealed a higher transcription of fructan-exohydrolases (FEH) in H11648 leaves (Fig. 4b) that may explain the differences found in monosaccharides content. The oligosaccharides fraction for *A. sisalana* leaves was also higher than the monosaccharide. In stem, *A. sisalana* presented higher oligosaccharides content than the other cultivars, when analyzing the transcriptome, we noticed that fructosyltransferases (FT) were more abundant in those samples.

#### 3.2.2. Compositional analysis of *Agave* leaves and stem

Regardless of the cultivar or tissue, the extractives represented the largest fraction of the biomass (Figure 6B). *A. fourcroydes* presented the highest content variation among its tissues, going from 31.2% in stem to 56.89% in leaf. The other cultivars and their tissues varied little, ranging from 49.3% to 52.94%. However, only *A. fourcroydes* stem was statically different from the other samples.

In general, the stems were more lignified than leaves. Considering leaves, *A. fourcroydes* lignin fraction was significatively lower than *A. sisalana* and H11648. For *A. fourcroydes,* lignin levels in leaves were practically half of those encountered for stem. Also, the stem of this agave cultivar presented the highest lignin content in all samples. No statistical difference was found between *A. sisalana* and H11648 lignin fractions. In relation to the cellulose content, *A. fourcroydes* stem presented the highest percentage (32.25%), with no statistical difference among the other samples. As for hemicellulose mass fraction, similar patterns to lignin were found. Leaves had lower hemicellulose contents and the stems, higher. There is no significant difference between the leaves of the cultivars. Nonetheless, among stems, hemicellulose fraction was significantly higher for *A. fourcroydes*, this cultivar presented 18.81%, which is 2.5x higher than its leaves. Concerning ashes, the highest values were found at *A. sisalana* leaves (11.28%), and the lowest was found at *A. fourcroydes* stem. In general, ash fractions in leaves were almost two times higher than stems.

#### 3.2.3. Enzymatic hydrolysis of cell wall carbohydrates

The hydrolysis protocol was effective in *Agave*, and, as expected, differences between samples were observed. Regardless of the agave cultivar, the stem showed higher recalcitrance than the leaves. The leaves of *A. fourcroydes* had the highest saccharification, while the stem of H11648 was the most recalcitrant sample. Compared to Energy Cane, *A. fourcroydes* leaves were 47% more hydrolysable, while *A. sisalana* and H11648 were 37%. Energy Cane hydrolysis was statistically equal to the stems of *A. sisalana* and *A. fourcroydes*. Within stems, *A. sisalana* and H11648 presented similar chemical composition; however, *A. sisalana* stem was the less recalcitrant. Interestingly, even with higher cellulose and lignin levels, the stem of *A. fourcroydes* was more accessible for digestion than the H11648 stem.

#### 3.2.4. Microscopic analysis of leaf anatomy and cell wall composition

The three cultivars presented thick cuticle, sunken stomata, and well-developed fiber caps, as previously described for *Agave* (Blunden, Yi, & Jewers, 1973). Pectic compounds deposition was mainly observed in the outer periclinal epidermal walls and mesophyll cells (supplementary material S4 a), as well as fibers around the vascular bundles, which stained intensely (figure 8 a-c). Compared to other cultivars, *A. sisalana* vascular bundle presented more fiber cap cells with thicker cell walls, which may explain the higher fiber quality of this cultivar (Medina, 1959). In contrast to pectin deposition, which was widespread, lignification was detected only in the secondary cell wall of the xylem conducting cells for all cultivars (figure 8 d-f). For the aniline blue staining, although we encountered brighter regions in the cuticle and fiber caps cells, which could indicate callose deposition, autofluorescence impaired the analysis (supplementary material S4 and S5).

**Figure 8.**
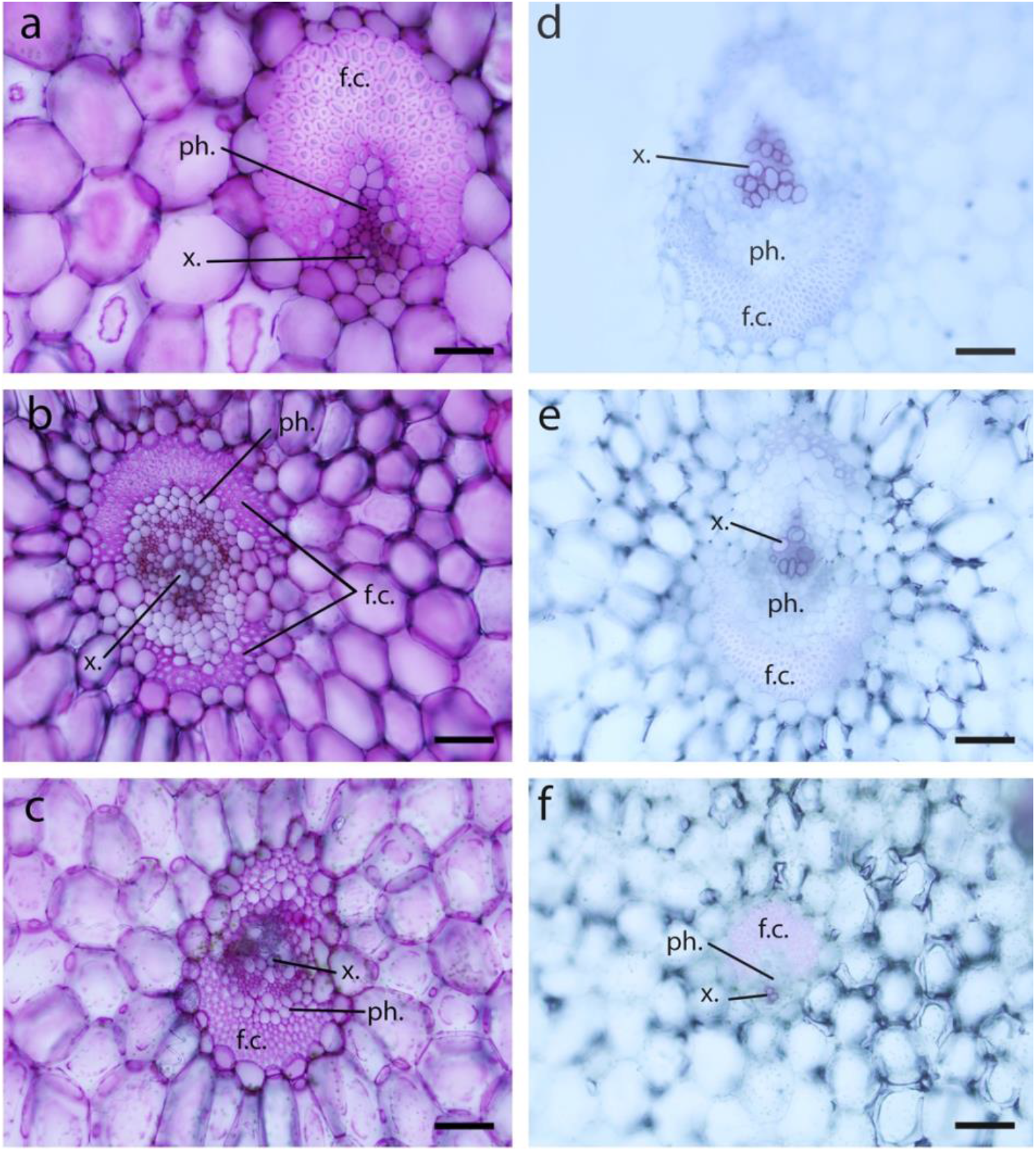
Lignin and Pectic compounds distribution in *Agave* vascular bundle. (a), (b), and (c) are leaf cross-sections from *A. sisalana*, H11648, and *A. fourcroydes*, respectively, stained with ruthenium red for pectic compounds detection; (d), (e), and (f) are leaf cross-sections stained with phloroglucinol-HCl for lignin detection of *A. sisalana*, H11648, and *A. fourcroydes*, respectively; (x.) xylem; (ph.) phloem; and (f.c.) fiber cap cells. Scale bars = 100 μm

## 4. DISCUSSION

Little is known about the chemical composition of agaves, especially for the cultivars utilized for fiber production. Few studies have been done on the subject, and most of the data available focus on the chemical composition of the fibers itself, not whole leaves or stems, which could be misleading when evaluating agave bioenergetic potential (Davis & Long, 2015; McDougall, Morrison, Stewart, Weyers, & Hillman, 1993; Mylsamy & Rajendran, 2010; Vieira, Heinze, Antonio-Cruz, & Mendoza-Martinez, 2002). Our data revealed some differences and common features between the fiber producing cultivars. In general, extractives represented a major part of the biomass composition, few differences were found on cellulose mass fraction, and agave stems were more lignified than leaves. Extractives, which are nonstructural components of biomass, may include pectin compounds and other waxes, phenolics, resin compounds, inorganic material, non-structural sugars, and nitrogenous material, among others (Pecha & Garcia-Perez, 2015). Using the same NREL’s normatives as we did, Yang *et al.*, 2015 has found a chemical composition in *A. tequilana* leaves similar to those reported here for the stems. The agaves developed for fiber production seem to have advantages from the industrial perspective considering the contents of lignin in leaves. Biorefineries based on agave leaves were suggested as an interesting alternative due to the possibility of maintaining a constant supply of plant material through the year, unlike stem-based industries in which the harvest occurs every 5-6 years (Davis, Dohleman, & Long, 2011; Davis & Long, 2015; Nobel, 2010; Yang et al., 2015). Compared to other crops, the lignin mass fraction, regardless of the tissue or cultivar, was lower than those found in woody biomass (21-32%, e.g., *Populus*, *Eucalyptus* and *Pinus*) (Ragauskas et al., 2014) and less or equal to found in herbaceous crops (9-18%, e.g., *Miscanthus*, *Panicum*, and corn straw) (Ragauskas et al., 2014).

In *Agave* leaves, lignin deposition occurs only on xylem vessels. Lignin is important for the xylem to hold its structure under negative pressure allowing water transport to occur properly (Campbell & Sederoff, 1996; Kitin et al., 2010). The lack of lignin in other leaf cells may be related to its hydrophobicity, as most of the agave leaf tissue is composed of water-storage cells of the mesophyll, and, in those cells, we encountered deposition of pectic compounds, which are hydrophilic. In addition, a high concentration of pectin in the cell wall was previously described to increase water absorption speed in leaves (Boanares et al., 2018), especially when accumulated in the epidermis. For all three cultivars, we encountered outer periclinal epidermis walls that are thicker with pectin deposition. The abundance of pectic compounds in leaves could contribute to the high percentage of extractives found in the compositional analysis.

Indeed, lignin content was the most interesting aspect of our chemical data, especially when analyzed together with the enzymatic hydrolysis assay. The differences observed on biomass accessibility could be explained by lignin mass fraction and possibly by its composition. The structure of lignin consists basically of three monomeric units: (I) syringyl (S); (II) guacyl (G); and (III) p-hydroxyphenyl (H) (Vanholme, Demedts, Morreel, Ralph, & Boerjan, 2010). The individual contribution of each monomer to the composition of this polymer varies significantly between tissues and species (Simmons et al., 2010). Higher rates of the S/G ratio are advantageous for the production of paper, cellulose and bioenergy, since the S-rich lignin is more easily dissociated from the cellulosic material (Lepikson-Neto et al., 2014; Simmons et al., 2010). It is possible that the differences found in the enzymatic hydrolysis of the stems, specifically the comparison between *A. fourcroydes* and H11648, are closely related to the S/G content. Although *A. fourcroydes* stem presented higher lignification, its biomass was less recalcitrant than H11648 stem. Moreover, the transcriptomic analysis of the phenylpropanoid pathway revealed that the expression of COMT, one of the main controlling points of the S lignin monomer biosynthesis, was higher in *A. fourcroydes,* suggesting that this cultivar is richer in S lignin, which would explain the higher accessibility of this tissue. The other recalcitrance oscillations may be partially explained by the differences in lignin mass fraction. In all cases, stems were more recalcitrant than leaves, and our expression data corroborate with this data, as most of the transcripts related to the lignin branch were more expressed in the stems.

Considering agave cell wall composition, it comes to one’s mind how can agaves be able to have such low lignin contents and still maintain their spatial structure supporting leaves that can achieve 1.7 meters long? Our phylogenetic analysis revealed an exclusive family of Callose synthase (CALS) between *Saccharum spontaneum*, *A. sisalana* and H11648. *S. spontaneum* was previously reported as having high expression and expanded families of CALS (Nascimento et al., 2019). Callose, a β-1,3-glucan polymer, is a cell wall component that is connected to a wide variety of plant processes, such as maintenance of the vascular system, plasmodesmata function, pollen and pollen tube development, cell plate formation, as well as biotic and abiotic defense responses (Falter et al., 2015; Schneider, Hanak, Persson, & Voigt, 2016). Although in our data, the exclusive CALS family did not present high expression, other CALS were differentially expressed and present at levels similar to those found for CsL, indicating that callose may play an important role in *Agave*. Also, low transcription of expansins were found, suggesting that the CALS transcription might not be connected to cell division and elongation activities. Other monocotyledonous energy crops with relativity low lignin content, such as maize and *Miscanthus x giganteus*, were found to form an unusual outer layer with callose fibrils interknitted in cellulose (Falter et al., 2015). In this context, we hypothesized that callose might be interacting with the agave cell wall and possibly thickening it to compensate for lower lignification through support and structure with the advantage to keep the cell wall elasticity, which is essential to allow the wide range of the water dynamics within the cells (filling and emptying in response to water availability). Fluorescence microscopy with aniline blue staining indicated a possible callose deposition at the fiber caps cells, which surrounds the vascular bundles, and are the main structural support for the leaves (Corbin et al., 2015; Rüggeberg et al., 2008). However, future analyzes are still required to confirm it. Perhaps, a novel strategy to genetically engineer crops with reduced lignin content without the typical decreased growth (Cesarino, Araújo, Domingues Júnior, & Mazzafera, 2013; Simmons et al., 2010; Vanholme, Ralph, et al., 2010) should consider improving callose fraction as well. Also, since callose is an easily degradable glucose polymer, it could still be harnessed for biofuels and biochemicals production.

However, low lignin contents do not come without negative collateral consequences, as positive correlations between lignin amount and pathogen resistance have been observed, especially for necrotrophic pathogens (Miedes, Vanholme, Boerjan, & Molina, 2014). Hence the importance of sisal bole rot disease caused by *Aspergillus welwitschiae*, a saprophytic fungus that infects sisal plants and behaves as a classical necrotrophic pathogen (Duarte et al., 2018). Many resistance mechanisms to opportunistic phytopathogens are related to plant secondary metabolites like flavonoids (Treutter, 2006). When analyzing the phenylpropanoid pathway, we were expecting that the flavonoid branch would present higher expression than the lignin one, since these two branches are competitive (Lepikson-Neto et al., 2014; X. Li, Bonawitz, Weng, & Chapple, 2010; Salazar et al., 2013). However, *CHS* transcript abundance was lower than *HCT*. Nevertheless, the phenylpropanoid pathway as a whole appears to be less expressed than other cell wall biosynthetic pathways such as cellulose. Precipitation, micrometeorological factors, and humidity are some of the most important components involved in the epidemiology of fungal pathogens and play a key role in both dispersion and spore germination (Bashi & Rotem, 1974; Buchanan, Gruissem, & Jones, 2015; Cook & Papendick, 1972; Huber & Gillespie, 1992; Rotem & Palti, 1969). It is possible that agaves can afford to express less of the phenylpropanoid pathway by inhabiting environments that are unfavorable to the development of fungal infections and, thus, redirecting valuable resources to abiotic stress resistance mechanisms.

In fact, agaves seem to invest heavily in drought and high-temperature resistance mechanisms. Our transcriptomic data revealed that the most expressed transcripts encode proteins that are well known to be stress-responsive. In *A. fourcroydes* leaves, *Phosphoenolpyruvate carboxylase*, responsible for the first step in CAM pathway, appears only in the fifth position of our most expressed transcripts list (Table 1) with an expression twenty-five times lower than the topmost (a Chaperone protein DnaJ); for *A. sisalana* this transcript is the fourth and for H11648 is the twenty-fourth. Since obligatory CAM species cannot count on transpirational cooling during the day and inhabit areas of high-light-intensity, these plants have developed alternative strategies to cope with high-temperature stress (Bita & Gerats, 2013; Borland et al., 2014; Sarwar et al., 2019). Our data suggest that for *Agave,* the molecular strategies may consist of overexpressing *HSP* and *LEA*, as well as genes related to proteolysis processes. These molecular mechanisms are present in all tissues regardless of the cultivar, yet, differences within cultivars were found at the transcript level. For instance, the most expressed *HSP* for *A. fourcroydes* was an *HSP40*, for *A. sisalana* it was an *HSP70*, and for H11648 a *small HSP*.

Considering LEA-encoding transcripts, the main difference appeared on expression levels between tissues. In general, *LEA5* was more abundant in leaves than stem and roots. Nonetheless, this transcript was one of the most expressed in every sample. Although their precise role has not been defined, LEA proteins may help prevent the formation of damaging protein aggregates due to desiccation or osmotic stresses (GOYAL, WALTON, & TUNNACLIFFE, 2005; Hundertmark & Hincha, 2008; Liu, Chakrabortee, Li, Zheng, & Tunnacliffe, 2011). It has been proposed that LEA proteins have different molecular mechanisms than chaperones, and evidence suggests that they can play a role as integral membrane protein (Caramelo & Iusem, 2009; Menze, Boswell, Toner, & Hand, 2009), and even protect mitochondrial membranes against dehydration damage (Caramelo & Iusem, 2009; Tolleter, Hincha, & Macherel, 2010). Also, previous work has correlated *LEA* expression with abiotic stress resistance in *Agave* (Tamayo-Ordóñez et al., 2016). Another interesting molecular strategy is proteolysis; we have found a GO term related to this process in leaves, and many ubiquitins highly expressed in all tissues and cultivars. Ubiquitylation has been reported to regulate many aspects of stress-response, being responsible for labeling damaged proteins for proteolysis by the proteasome system (Flick & Kaiser, 2012a). Thereby, the cells can remove proteins damaged by stress that are unable to function properly, preventing the accumulation of potentially harmful aggregates and recycling nutrients, especially nitrogen (Flick & Kaiser, 2012a; Lyzenga & Stone, 2012).

Many plants accumulate compatible osmolytes to endure drought and high salinity stress through osmoprotection, but these compounds vary among species. For example, while the presence of the amino acid proline as a compatible solute is widespread in plants, just a few members of *Plumbaginaceae* family accumulates β-alanine betaine as an osmolyte (Buchanan et al., 2015; Rathinasabapathi, Fouad, & Sigua, 2001). One important osmolyte is trehalose, whose biosynthetic genes were differentially expressed, as well as an unshared family in *A. sisalana.* Previous transcriptomic data have suggested that, in *Agave sisalana*, trehalose may be an osmolyte (Sarwar et al., 2019). However, in our samples, no actual trehalose was detected by the gas chromatography. Instead, our chemical analysis detected the raffinose in every sample tested. We suggest that, in *Agave*, one of the main osmolytes might be raffinose. This carbohydrate was detected insimilar concentration in every sample tested, yet our transcriptomic data revealed clear distinction regarding the raffinose biosynthetic pathways; whereas *A. fourcroydes* seems to prefer the RafS-mediated synthesis, the other two cultivars seem to utilize the route mediated by GosL. It is plausible that the inclination of *A. fourcroydes* to use RafS-mediated synthesis could affect cellulose biosynthesis since SUSY and RafS could compete for the same substrate.

Even though leaves presented higher expression of genes related to abiotic stresses, it was in roots that we have found many of the GO terms referred to these responses (e.g., response to oxidative stress; response to stress). Soil surface can have large daily oscillations in temperature, and in desert-like regions, these temperatures can easily overcome 40°C (Nobel, 2010; Sattari, Dodangeh, & Abraham, 2017). As agaves present shallow roots systems (Nobel, 1988), this tissue is continuously exposed to harsh conditions and must develop special adaptations to cope with high temperature, salinity, and water deficit. It makes sense that the most abundant transcripts of the root-associated fungi are of proteins related to heat responses, which may indicate that the microbial community is well adapted to such an environment.

## 5. CONCLUSION

Drought is one of the most important environmental factors that impact plant growth and development. In the eminence of climate change, projections estimate disturbances in rainfall patterns and rise in global temperature, arid and semi-arid environments might become more common, and plants will have to be adapted to those conditions. Therefore, understanding relevant agronomical examples at molecular level can be the key to the development of new technologies. Perhaps, *Agave* may offer the ideal conditions do decipher the blueprint for biomass production under dry, hot conditions. Our inspection of these plants, thriving under extreme conditions, indicated that abiotic stress mechanisms pervade this crop metabolism. Stress-response genes were the most highly expressed transcripts, and, also, many of the plant processes have secondary functions that act in response to stress, from reserve carbohydrates that can stabilize membranes and protect cells against the effects of dehydration (Livingston, Hincha, & Heyer, 2009), photosynthetic mechanisms which are highly water-use efficient (Stewart, 2015), to cell wall components and adaptations that can control water flux (Boanares et al., 2018; De Storme & Geelen, 2014). It is possible that the fine-tuning between these auxiliary functions and the main resistance mechanisms is what makes agaves so adapted to marginal ecosystems and still great biomass producers. In summary, our data constitute an important new resource for the study of *Agave* species and indicate potential mechanisms that could be used to improve the tolerance of other crops to drought and high-temperature conditions.

## Supporting information

Table S1 Monthly precipitation (mm) of Monteiro-PB from January 1995 to July 2016

Table S2 List of all the assembled transcripts with corresponding annotation, quantification (TPM), and specificity measure (SPM)

Table S3 List of the gene families generated by ORTHOMLC and ortholog groups of the comparative genomics analysis

## 6. ACKNOWLEDGEMENTS

This work was supported in part by the *Coordenação de Aperfeiçoamento de Pessoal de Nível Superior* - Brazil (CAPES) - Finance Code 001, Center for Computational Engineering and Sciences - FAPESP/Cepid (2013/08293-7), and *Fundação de Amparo à Pesquisa do Estado de São Paulo* - Brazil (FAPESP) - process numbers 2016/05396-8 and 2017/04900-7.

