## Supplementary Information for "Transcriptome analysis of three *Agave* fiber-producing cultivars suitable for biochemicals and biofuels production in semiarid regions"

**Table S1** Monthly precipitation (mm) of Monteiro-PB from January 1995 to July 2016. Data were retrieved from the BDMEP database *(http://www.inmet.gov.br/projetos/rede/pesquisa/)* of the National Institute of Meteorology of Brazil. (Excel file separately attached).

**Table S2** List of all the assembled transcripts with corresponding annotation, quantification (TPM), and specificity measure (SPM) (Excel file separately attached).

**Table S3** List of the gene families generated by ORTHOMLC and ortholog groups of the comparative genomics analysis (Excel file separately attached). AF: A. fourcroydes, AS: A. sisalana, HY: H11648.

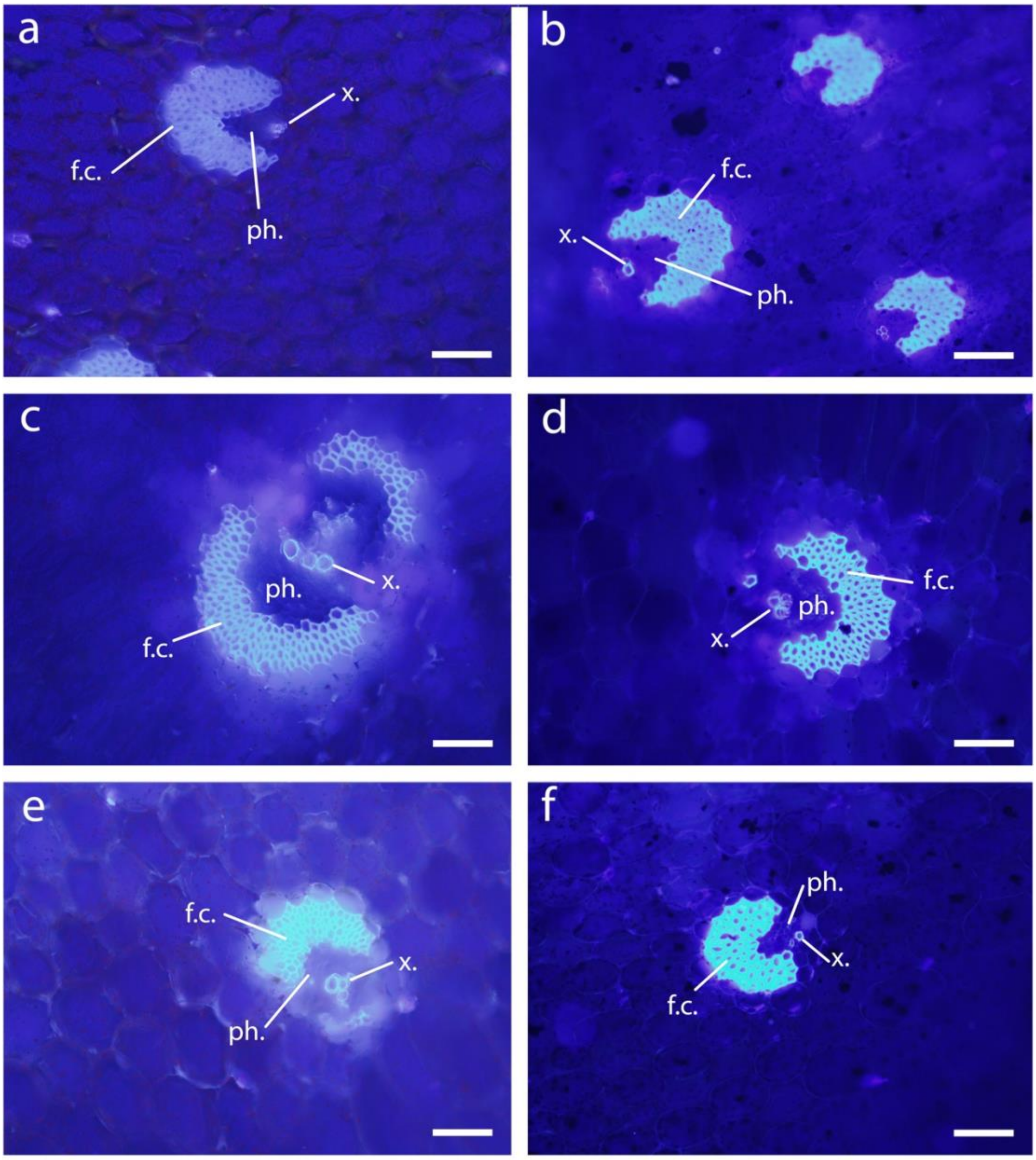


**Figure S4** Fluorescence microscopy with aniline blue staining in *Agave* vascular bundle. (a), (c), and (e) are leaf cross-sections from *A. sisalana*, H11648, and *A. fourcroydes*, respectively, without aniline blue staining and observed under an epifluorescence microscope with a UV filter (BP 340 to 380 nm, LP 425 nm); (b), (d), and (f) are leaf cross-sections stained with aniline blue of *A. sisalana*, H11648, and *A. fourcroydes*, respectively, and observed under the same microscopy conditions. (x.) xylem; (ph.) phloem; and (f.c.) fiber cap cells. Scale bars = 100 µm.

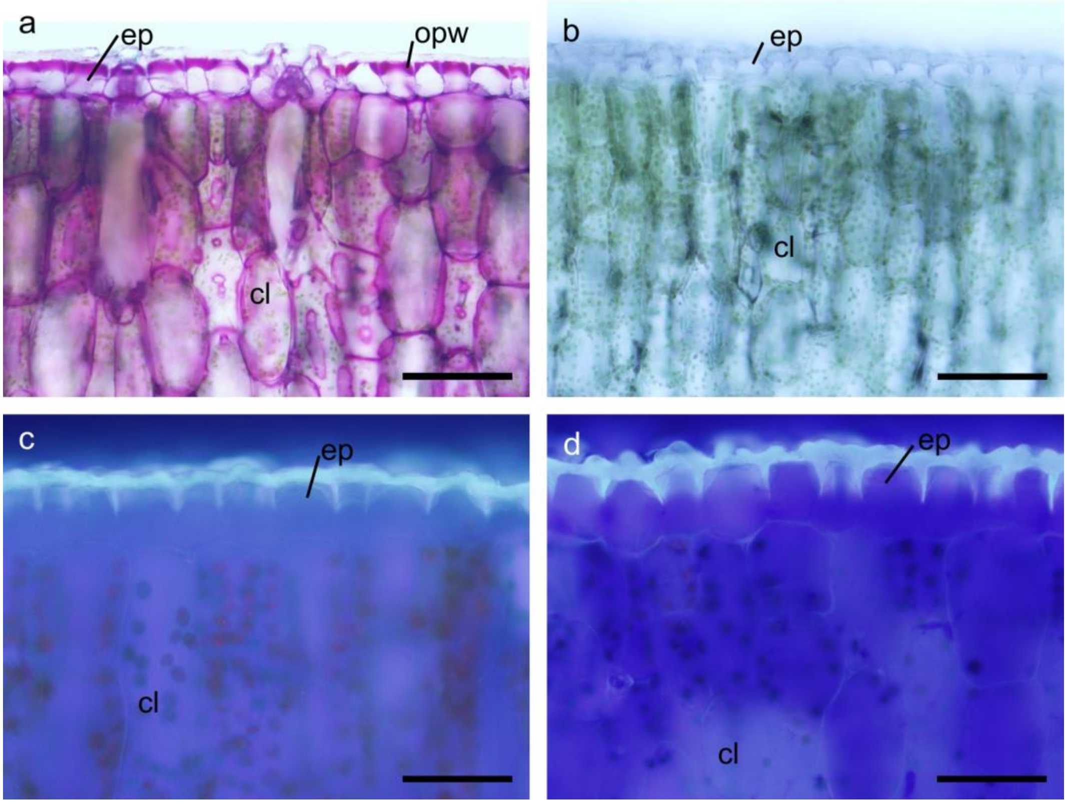


**Figure S5** Cross-sections from *Agave* leaf epidermis. (a) e (b) *A. sisalana*; (a) stained with ruthenium red for pectic compounds detection, detail of the thick outer periclinal walls with pectin deposition evidenced in pink; (b) leaf cross-section stained with phloroglucinol-HCl for lignin detection; (c) e (d) H11648; (c) without aniline blue staining and observed under an epifluorescence microscope with a UV filter; (d) with aniline blue staining and observed under an epifluorescence microscope with a UV filter. (cl) chlorenchyma; (ep) epidermis; (opw) outer periclinal walls. Scale bars = 50 µm.
